# Atypical small GTPase RABL3 interacts with RAB11 to regulate early ciliogenesis in human cells

**DOI:** 10.1101/2022.03.14.484356

**Authors:** Tetsuo Kobayashi, Tatsuya Ikeda, Reo Ota, Takafumi Yasukawa, Hiroshi Itoh

## Abstract

Primary cilia are near-ubiquitously assembled on cells in the human body and are broadly associated with genetic diseases and cancers. In the early stage of ciliogenesis, the ciliary vesicle (CV) is formed on the mother centriole, which nucleates the primary cilium. However, the regulatory mechanisms underlying CV formation have not yet been fully elucidated. Here, we found that the atypical small GTPase RAB-Like 3 (RABL3) is necessary to assemble primary cilia in human cells. RABL3 directly interacts with RAB11, which is involved in CV formation. RABL3 localizes around the centrosome during early ciliogenesis, reminiscent of RAB11 dynamics. Furthermore, RABL3 positively controls the CV formation like RAB11. These findings suggest that RABL3 plays an important role, in cooperation with RAB11, in CV formation during early ciliogenesis.

## Introduction

The centrosome contains two cylinder-like structures, termed mother and daughter centrioles. The mother centriole is equipped with distal and subdistal appendages (DA and SDA, respectively) that are absent from the daughter centriole. Centrioles nucleate spindles during mitosis, whereas the mother centriole becomes the base of a sensory organelle called the primary cilium during interphase (Kobayashi and Dynlacht, 2011; Sánchez and Dynlacht, 2016). The primary cilium protrudes from the cell surface of most mammalian cells and contains multiple signaling molecules (Ishikawa and Marshall, 2011). Structural and functional anomalies in primary cilia are implicated in a broad spectrum of genetic diseases and cancers (Eguether and Hahne, 2018; Liu et al., 2018; Reiter and Leroux, 2017). In cultured mammalian cells, serum deprivation from the culture medium induces a transition from the mother centriole to the primary cilium. A small vesicle, termed ciliary vesicle (CV), covers the top of the mother centriole in the early stage of the intracellular ciliogenesis in human cells (SOROKIN, 1962).

Pre-ciliary vesicles (PCVs), which are thought to be derived from the Golgi, are initially attached to the DA of the mother centriole, where they are referred to as distal appendage vesicles (DAVs), and DAVs assemble into the CV (Shakya and Westlake, 2021). The CV extends along with microtubule elongation and finally develops into the ciliary membrane (CM), encapsulating the ciliary axoneme. Thus far, mounting studies have indicated that several proteins regulate CM formation processes in a stepwise manner, including the RAB-family small GTPases. Myo-Va occupies the PCV and aids in the trafficking of PCVs to the DA (Wu et al., 2018). EHD1/3, SNAP29, PACSIN1/2, and MICAL-L1 mediate CV assembly from DAVs (Insinna et al., 2019; Lu et al., 2015). The small GTPase RAB34 is also required for CV assembly step (Ganga et al., 2021; Stuck et al., 2021). The small GTPase RAB8 appears to be involved in the extension of CV to develop CM (Lu et al., 2015). Two ciliopathy-related proteins, Talpid3 and CEP290, are necessary for RAB8 recruitment and proper CV development (Kobayashi et al., 2014). RAB8 is activated by a guanine nucleotide exchange factor (GEF), Rabin8, and the small GTPase RAB11 (Knödler et al., 2010). RAB11 regulates GEF activity and centrosome targeting of Rabin8 (Knödler et al., 2010; Westlake et al., 2011). Recently, it was shown that lysophosphatidic acid (LPA)/LPA Receptor-1-dependent PI3K/Akt signaling controls RAB11 activation during ciliogenesis (Walia et al., 2019). RAB-Like 3 (RABL3) is a member of the RABL family, comprising a RAB GTPases subfamily (Colicelli, 2004). RABL proteins are devoid of sites for lipid modification and some conserved residues in GTP-binding proteins (Blacque et al., 2018; Homma et al., 2021). RABL3 is upregulated in several cancers and is required for their proliferation and migration (An et al., 2017; Ge et al., 2019; Li et al., 2010; Ma et al., 2021; Pan et al., 2017; Usman et al., 2020; Xu et al., 2021; Zhang et al., 2016). Mutations in *rabl3* are correlated with heritable pancreatic cancer (Nissim et al., 2019). A recent study showed that RABL3 is crucial for lymphoid function, and that its knockout mice are embryonic lethal (Zhong et al., 2020). Interestingly, four of the six RABL proteins are known to be involved in molecular trafficking to and within primary cilia. RABL4/IFT27 and RABL5/IFT22 comprise the intraflagellar transport (IFT) machinery, a universally conserved protein complex that bi-directionally conveys molecules in primary cilia (Nakayama and Katoh, 2018). RABL2 controls anterograde IFT and trafficking of ciliary G-protein-coupled receptors (Dateyama et al., 2019; Kanie et al., 2017; Nishijima et al., 2017). A recent study showed that RABL2 ensures export of ciliary proteins (Duan et al., 2021). A proteomic analysis using mouse photoreceptor sensory cilia suggested RABL3 as a cilia-related protein (Liu et al., 2007); however, it is unknown whether and how RABL3 is associated with primary cilia.

In this study, we investigated normal diploid human cells depleted of RABL3 and discovered that RABL3 is required to assemble primary cilia. A proteomic approach identified RAB11 as a RABL3-binding protein. RABL3 directly interacts with and is stabilized by RAB11 *in vitro*. RABL3 accumulated around the centrosome during early ciliogenesis, similar to RAB11. Furthermore, RABL3 depletion impaired CV formation, but ectopic RABL3 expression promoted CV formation, phenocopying ablation and overexpression of RAB11, respectively. Altogether, these results suggest that RABL3 cooperates with RAB11 and regulates the CV formation during early ciliogenesis in human cells.

## Results

### RABL3 is required for primary ciliogenesis in RPE1 cells

To investigate the consequences of RABL3 depletion in normal diploid human cells, we generated RABL3-mutated retinal pigment epithelial (RPE1) cells by CRISPR/Cas9-mediated gene editing. Sequence analysis indicated heterozygous mutations in two clones, Rabl3-1 and Rabl3-2 (Figure S1A). While a four-nucleotide deletion in one allele led to a premature stop codon before the G4 domain in Rabl3-1, a three-nucleotide in-frame deletion resulting in replacement of two amino-acids, (glutamine and asparagine) with one amino-acid (histidine) occurred in another allele. This QN motif is conserved in vertebrate RABL3 (Figure S1B), suggesting that this mutation seriously impairs RABL3. In contrast, mutations in both alleles lead to premature stop codons in Rabl3-2. We found a substantial decrease in RABL3 protein levels in both clones by immunoblotting analysis, although RABL3 was faintly detected in Rabl3-1 as predicted by sequence analysis (Figure 1A).

**Figure 1.**
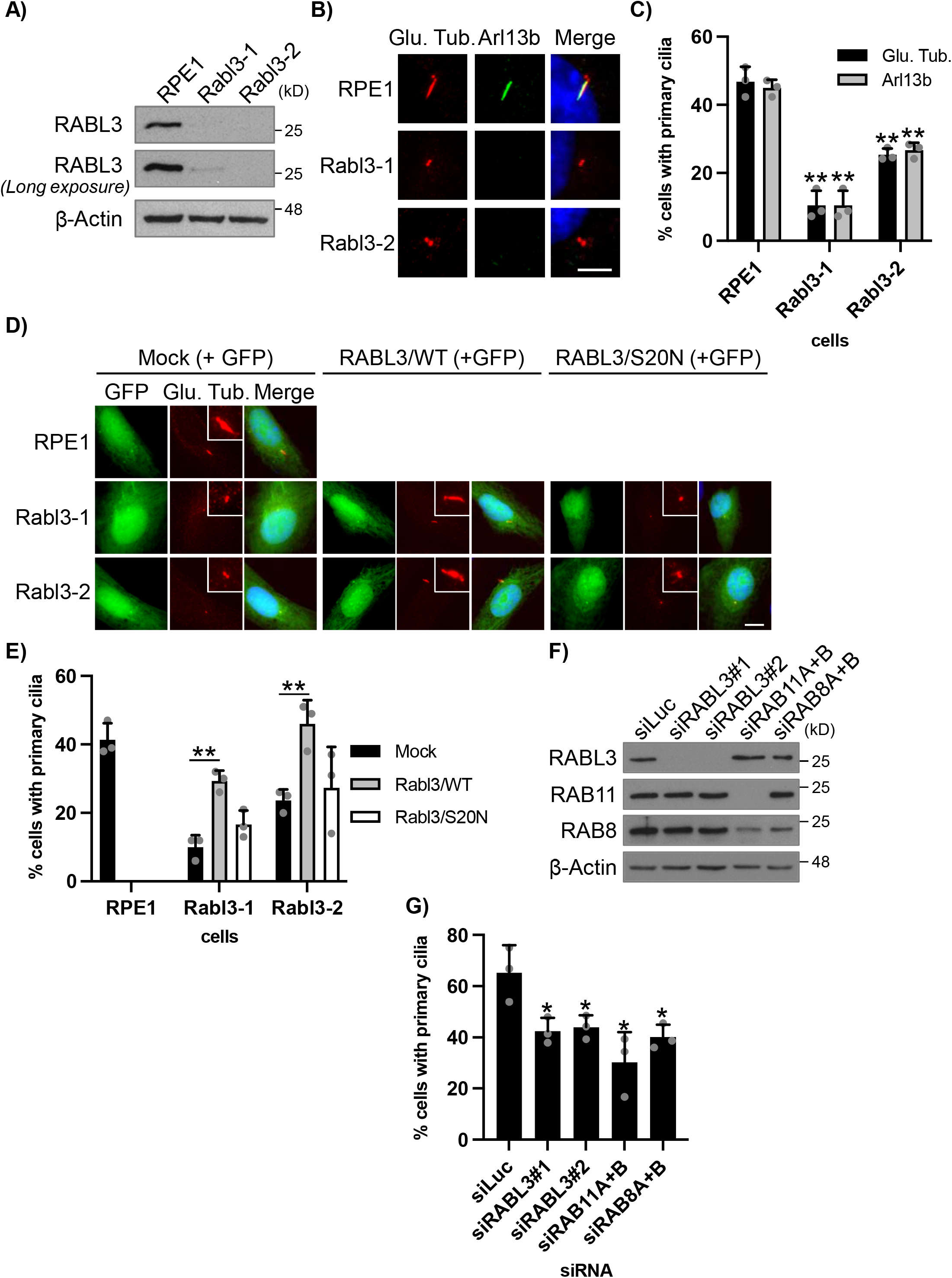
RABL3 depletion impairs primary cilia formation in RPE1 cells. **(A)**Indicated cells were cultured for 48 hrs. Cell extracts were immunoblotted with an anti-RABL3 antibody. β-Actin was used as a loading control. **(B, C)** Indicated cells were cultured in serum-starved medium for 24 hrs and immunostained with anti-glutamylated tubulin (GT335, red) and anti-Arl13b (green) antibodies. (B) DNA was stained with Hoechst (blue). Scale bar, 5 μm. (C) The percentage of cells with primary cilia was determined. Average of three independent experiments is shown. >250 cells were scored for each experiment. **(D, E)** Indicated cells transfected with plasmids expressing EGFP and mock, Flag-hRABL3/WT, or S20N were cultured in serum-starved medium for 24 hrs and immunostained with an anti-glutamylated tubulin antibody (red). (D) DNA was stained with Hoechst (blue). Scale bar, 5 μm. (E) The percentage of GFP-positive cells with primary cilia was determined. Average of three independent experiments is shown. >100 cells were scored for each experiment. **(F, G)** RPE1 cells transfected with indicated siRNA were cultured in serum-starved medium for 48 hrs. (F) Cell extracts were immunoblotted with indicated antibodies. β-Actin was used as a loading control. (G) Cells were immunostained with an anti-glutamylated tubulin antibody. The percentage of cells with primary cilia was determined. Average of three independent experiments is shown. >250 cells were scored for each experiment. **(C, E, G)** Error bars represent SD. *: *p* < 0.05, **: *p* < 0.01 compared with RPE1 (A), Mock (E), or siLuc (F).

We then investigated primary cilia formation in RABL3-mutated cells. Primary cilia were visualized by immunofluorescence experiments with two specific antibodies against glutamylated tubulin (Glu. Tub.) and ARL13B. The wild-type (WT) RPE1 cells assembled primary cilia when induced to quiescence by depriving the serum in the culture medium (Figure 1B, C). In contrast, Rabl3-1 and Rabl3-2 cells formed significantly fewer primary cilia than WT cells (Figure 1B, C). We next conducted rescue experiments to verify that the de-ciliation phenotype in RABL3-mutated cells was due to the loss of RABL3. Ectopic expression of RABL3 significantly restored primary cilia in Rabl3-1 and Rabl3-2 cells (Figure 1D, E). In contrast, the RABL3/S20N mutant, which is expected to fail to bind GTP, namely a negatively locked mutant, did not ameliorate the phenotype (Figure 1D, E). These results suggest that RABL3 with GTP-binding capacity is required for ciliogenesis. We also performed siRNA-mediated knockdown experiments. The protein expression of endogenous RABL3 was considerably decreased by introducing two individual siRNAs targeting RABL3 (Figure 1F). We observed a significant reduction in primary cilia assembly in these RABL3-ablated RPE1 cells (Figure 1G). Collectively, these results strongly suggest that RABL3 is a requisite for primary cilia formation in RPE1 cells.

### RABL3 interacts with and is stabilized by RAB11

To uncover the mechanistic role of RABL3 in primary ciliogenesis, we performed a proteomic screen to identify RABL3-interacting proteins. Immunoaffinity chromatography and subsequent mass spectrometric analysis identified a peptide common to two close paralogs of small GTPases, RAB11A and RAB11B, in RABL3 immunoprecipitation (Figure S2A, B). We co-expressed Flag-tagged RABL3 and GFP-fused RAB11A in HEK293T cells and performed anti-Flag immunoprecipitation to test if they interact in cells and found that RABL3 co-precipitated with RAB11A (Figure 2A). We subsequently performed similar experiments using cells co-expressing RABL3/WT or S20N, and RAB11A/WT, actively locked Q70L mutant, or negatively locked S25N mutant, and found efficient co-precipitation of RABL3/S20N and RAB11A/Q70L (Figure 2A). Reciprocal immunoprecipitation using an anti-GFP antibody confirmed similar binding properties (Figure 2B). These results suggest that the negative form of RABL3 preferentially interacts with the active form of RAB11A. We next investigated the direct interaction between RABL3 and RAB11. To this end, we purified recombinant RABL3 and RAB11A proteins from bacterial lysates (Figure 2C). We then performed a pull-down assay using glutathione sepharose with buffer containing Mg^2+^, and found that GST-RAB11A specifically pulled down RABL3 (Figure 2D, left), demonstrating their direct binding. In contrast, adding a non-hydrolyzable GTP analog GTPγS or GDP substantially abrogated their association (Figure 2C, middle and right). This result shows that RABL3 and RAB11 fail to bind to each other in the same guanine nucleotide forms, consistent with previous immunoprecipitation assays that the negative form of RABL3 associated with the active form of RAB11A.

**Figure 2.**
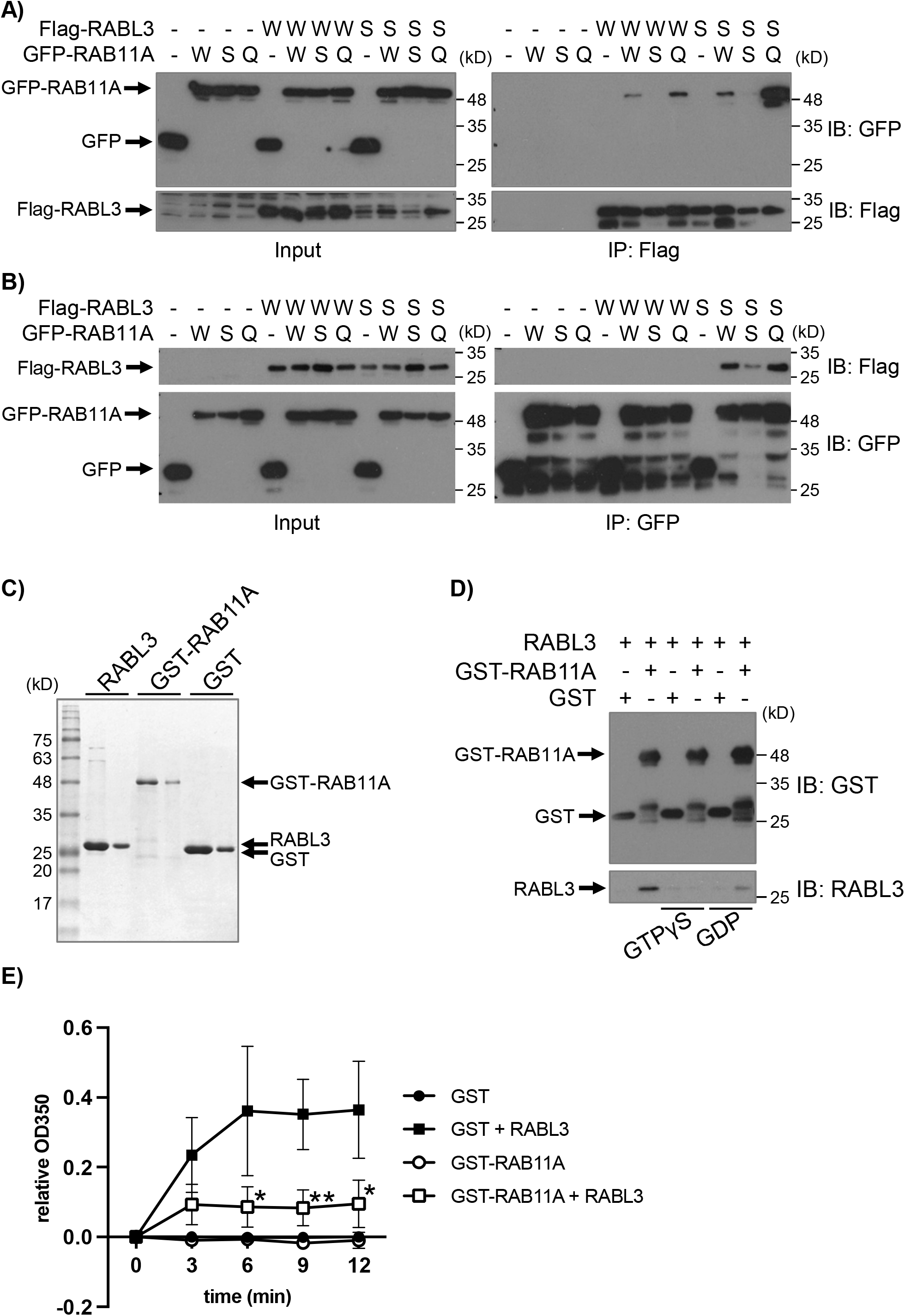
RAB11 interacts with and stabilizes RABL3. **(A, B)** Flag-RABL3/WT (W) or S20N (S) and GFP-RAB11A/WT (W), Q70L (Q) or S25N (S) were expressed in HEK293T cells. Lysates were immunoprecipitated with (A) anti-Flag or (B) anti-GFP (GFP-nanobody) antibodies. The resulting immunoprecipitates were western blotted with indicated antibodies. **(C)** Purified recombinant proteins visualized by Coomassie stain. **(D)** Recombinant RABL3, GST-RAB11 and GST proteins were subjected to *in vitro* binding assay in the presence of indicated additives. The resulting precipitates were immunoblotted with indicated antibodies. **(E)** Recombinant RABL3, GST-RAB11 and GST were subjected to Turbidity assay. The optimal density at 350 nm of the protein solution was determined every 3 mins. Average of three independent experiments is shown. Error bars represent SD. *: *p* < 0.05, **: *p* < 0.01 compared with GST + RABL3.

A previous study showed that RABL4/IFT27 binds to the small GTPase ARL6 and prevents the aggregation of ARL6 *in vitro* (Liew et al., 2014). Because of the analogous RABL-small GTPase interaction, we performed a similar turbidity assay using the recombinant RABL3 and RAB11A proteins to test mutual chaperone activity. The optical density at 350 nm was monitored to detect the insoluble (precipitated) proteins (Liew et al., 2014). While RABL3 was precipitated by incubation at 37 °C for 12 min, the precipitation was significantly reduced in the mixture of RABL3 and GST-RAB11A (Figure 2E). In contrast, GST-RAB11A was soluble in this assay. These results suggest that RAB11 stabilizes RABL3 *in vitro*.

### RABL3 accumulates around the centrosome during early ciliogenesis

We assessed the subcellular localization of RABL3 in RPE1 cells by performing immunofluorescence assays. Multiple puncta were detected throughout the cells with a RABL3 antibody (Figure 3A, upper left), and these puncta were almost completely absent in Rabl3-2 cells (Figure 3A, lower left), indicating that the RABL3 antibody specifically recognizes endogenous RABL3 in RPE1 cells. Endogenous RABL3 incrementally accumulated in the vicinity of the centrosome during induction to quiescence for 6 h in RPE1 cells but not in Rabl3-2 cells (Figure 3A, right, 3B). These RABL3-positive puncta around CEP164-positive mother centriole partially overlapped with RAB11, which is known to accumulate around the centrosome during early ciliogenesis (Figure 3C) (Westlake et al., 2011). In addition, a fraction of RABL3 co-localized with RAB11 and GM130, a canonical marker of the Golgi, after 6 h of serum-starvation (Figure 3D). These results collectively indicate that RABL3 is partly located around the centrosome and the Golgi apparatus and co-localizes with RAB11 during early ciliogenesis. Furthermore, to evaluate guanine nucleotide-form-dependent localization of RABL3 during ciliogenesis, RPE1 cells expressing Myc-tagged RABL3/WT or S20N were cultured in the serum-starved medium for 6 h. Although RABL3/WT was distributed uniformly or marginally around the centrosome, RABL3/S20N was clearly concentrated at the centrosome (Figure 3E, right). These observations suggest that the negative form of RABL3 is recruited to the centrosome during early ciliogenesis.

**Figure 3.**
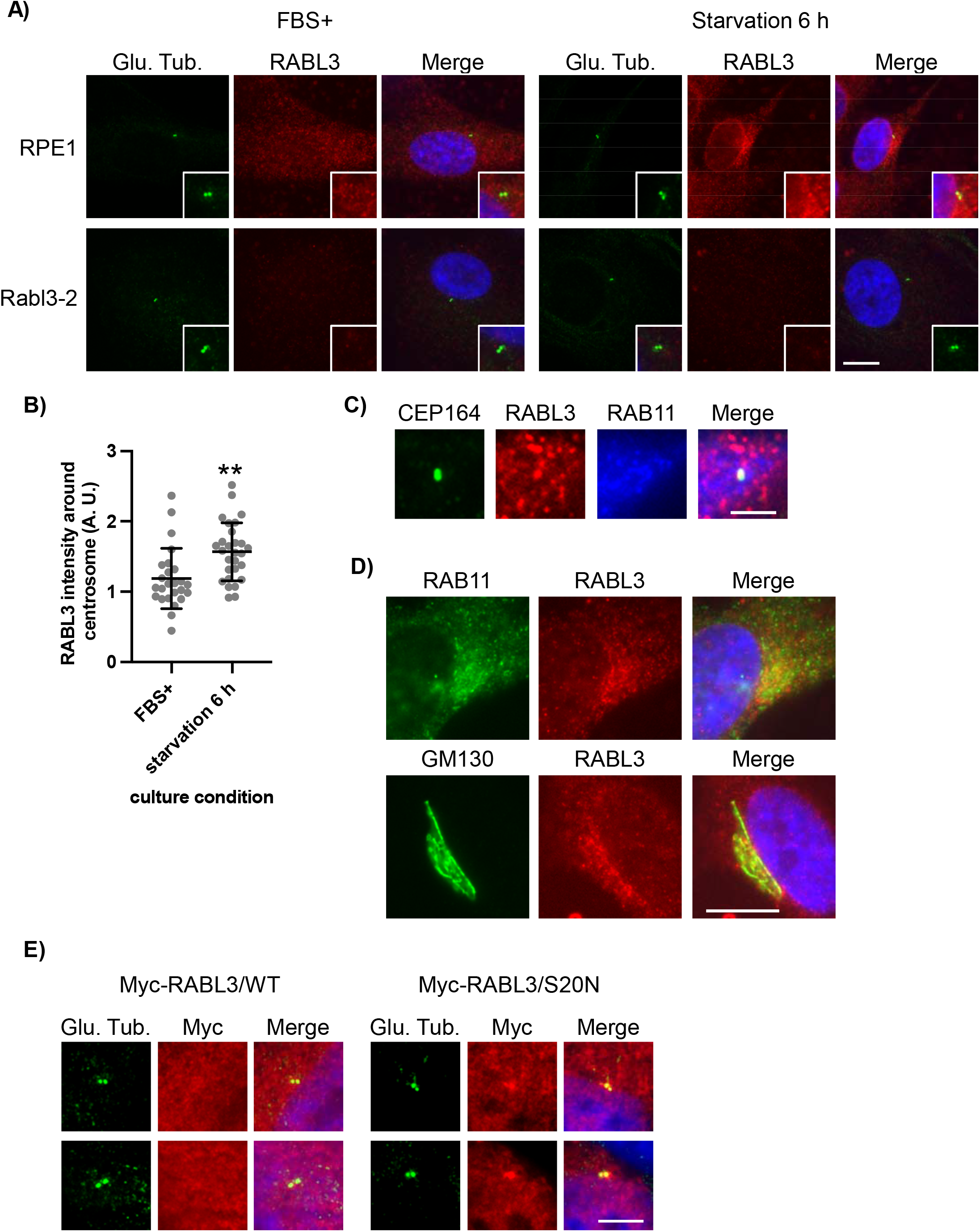
RABL3 accumulates around centrosomes during early ciliogenesis. **(A, B)** Indicated cells were cultured in serum-fed medium for 48 hrs (left) or serum-starved medium for 6 hrs (right) and immunostained with anti-glutamylated tubulin (green) and anti-RABL3 (red) antibodies. (A) DNA was stained with Hoechst (blue). Scale bar, 10 μm. (B) The quantified fluorescence intensity of RABL3 within ∼3 μm diameter circle around centrosome is shown. n = 25 (FBS +), 28 (Starvation 6 h). **(C)** RPE1 cells were cultured in serum-starved medium for 6 hrs and immunostained with anti-CEP164 (green), anti-RABL3 (red), and anti-RAB11 (blue) antibodies. Scale bar, 2.5 μm. **(D, E)** RPE1 cells were cultured in serum-starved medium for 6 hrs and immunostained with anti-RABL3 (red) and (D) anti-RAB11 (green) or (E) anti-GM130 (green) antibodies. DNA was stained with Hoechst (blue). Scale bar, 10 μm. **(E)** RPE1 cells transfected with plasmids expressing Myc-hRABL3/WT (left) or S20N (right) were cultured in serum-starved medium for 6 hrs and immunostained with anti-glutamylated tubulin (green) and anti-Myc (red) antibodies. Two representative images are shown. DNA was stained with Hoechst (blue). Scale bar, 5 μm. **(B)** Error bars represent SD. **: *p* < 0.01 compared with FBS +.

### RABL3 depletion impedes early ciliogenesis

We attempted to reveal the role of RABL3 in the primary cilium formation process. We first examined CP110, which localizes to both mother and daughter centrioles in cycling cells and disappears from the mother centriole during early ciliogenesis (Spektor et al., 2007). We found that cells with two CP110 dots were significantly increased upon silencing RABL3 (Figure 4A, B), suggesting that it is required for the CP110 removal from the mother centriole. RAB11 depletion also hampered primary ciliogenesis and the loss of CP110 dots (Figure 1F, G, 4A, B). Next, we explored GFP-Rabin8 vesicles that accumulate around the centrosome immediately after serum withdrawal in RPE1 cells (Westlake et al., 2011). Ablation of RABL3 suppressed the accumulation of GFP-Rabin8 around the centrosome (Figure 4C, D). RAB11 knockdown induced a similar phenotype (Figure 4C, D), consistent with previous reports (Lu et al., 2015; Westlake et al., 2011). These results suggest that RABL3 is a prerequisite for the early steps of primary cilia formation, similar to RAB11.

**Figure 4.**
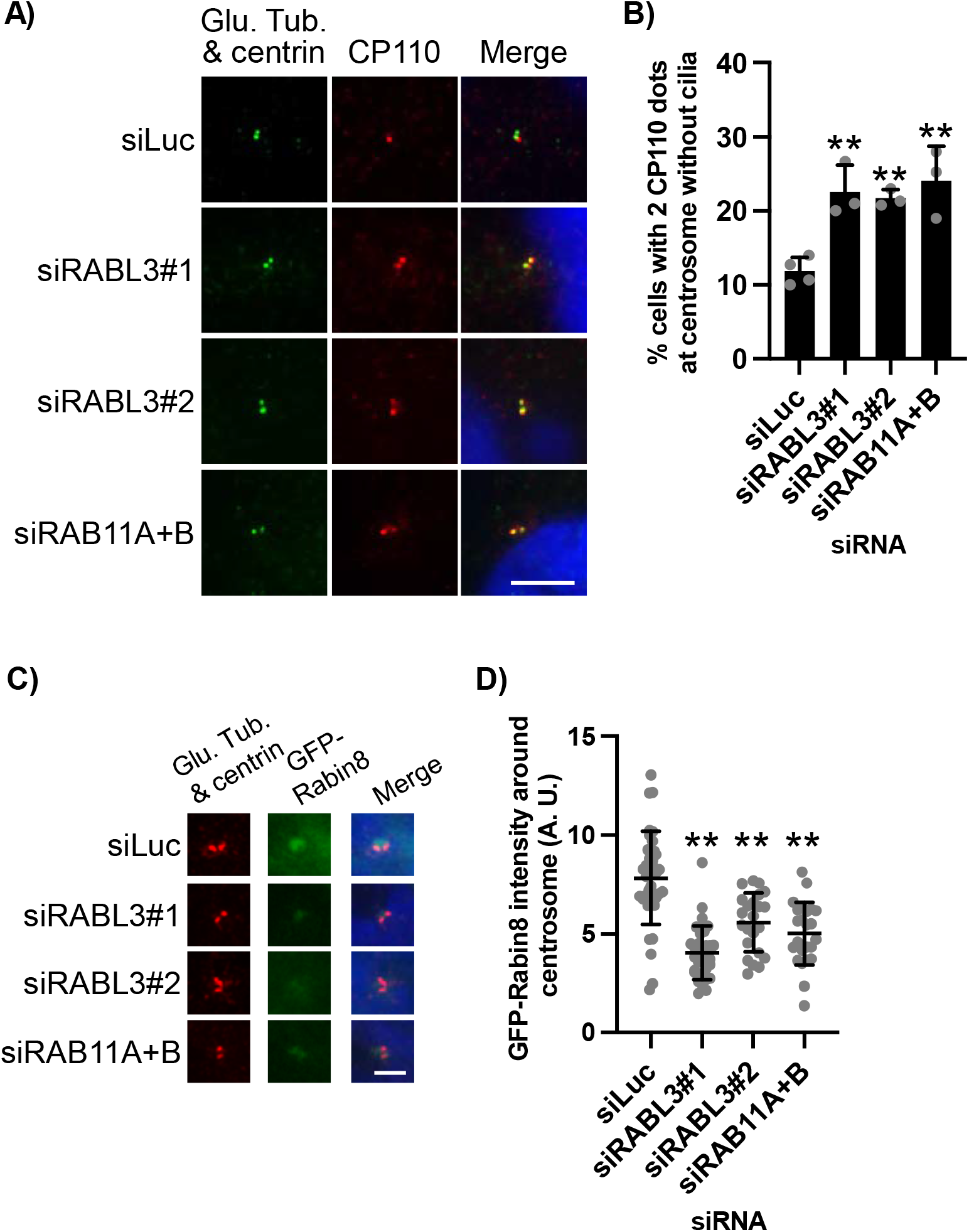
RABL3 is required for early ciliogenesis. **(A, B)** RPE1 cells transiently transfected with indicated siRNA were cultured in serum-starved medium for 48 hrs. Cells were immunostained with anti-glutamylated tubulin (green), anti-centrin (green), and anti-CP110 (red) antibodies. (A) DNA was stained with Hoechst (blue). Scale bar, 5 μm. (B) The percentage of cells with two CP110 dots at non-ciliated centrosome was determined. Average of three to four independent experiments is shown. >100 cells were scored for each experiment. **(C, D)** GFP-Rabin8-RPE1 cells transiently transfected with indicated siRNA were cultured in serum-starved medium for 1 hr. Cells were immunostained with anti-glutamylated tubulin (red) and anti-centrin (red) antibodies. (C) DNA was stained with Hoechst (blue). Scale bar, 2.5 μm. (D) The quantified fluorescence intensity of GFP-Rabin8 within ∼3 μm diameter circle around non-ciliated centrosome is shown. n = 37 (siLuc), 32 (siRABL3#1), 23 (siRABL3#2), 24 (siRAB11A+B). **(B, D)** Error bars represent SD. **: *p* < 0.01 compared with siLuc (B, D).

### RABL3 positively controls CV formation

As GFP-Rabin8 vesicles are thought to provide membrane components to assemble ciliary membrane structures (CV and CM) (Shakya and Westlake, 2021), we established GFP-Smoothened (SMO)-expressing RPE1 cells to visualize their formation (Lu et al., 2015). While elongated GFP-SMO along with the axoneme, indicating CM, was observed in control cells, RABL3 or RAB11 depletion abolished GFP-SMO-positive foci at the centriole (Figure 5A, B), suggesting that RABL3 and RAB11 are required for GFP-SMO-positive CV (SMO^+^ CV) formation. In contrast to RABL3 and RAB11, silencing of RAB8 allowed SMO^+^ CV formation irrespective of suppressed ciliation (Figure 1F, G, 5A, B). These data suggest that RABL3 and RAB11 play overlapping roles in CV formation, whereas RAB8 is necessary after CV formation, as described previously (Lu et al., 2015).

**Figure 5.**
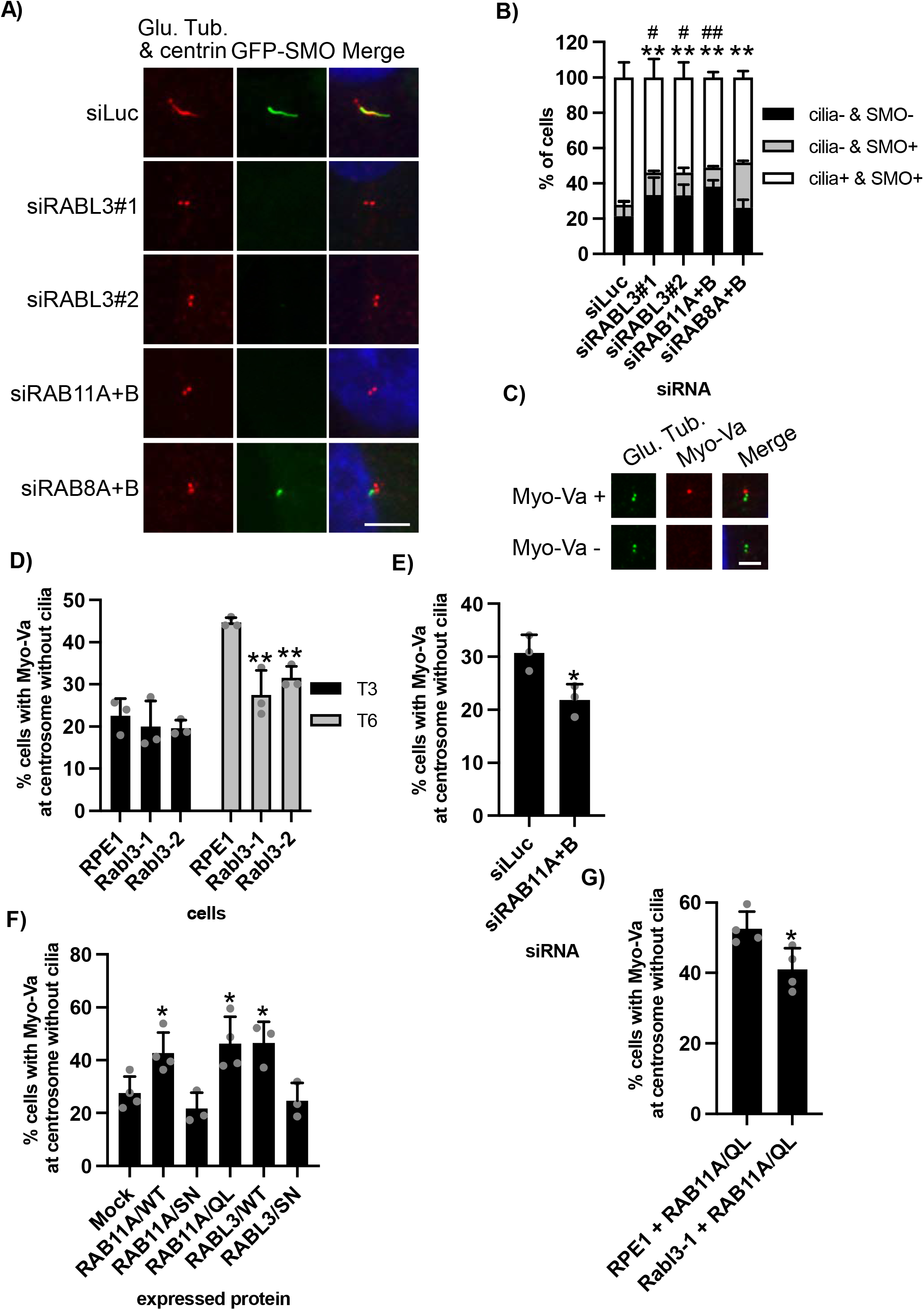
RABL3 positively controls CV formation like RAB11. **(A, B)** GFP-SMO-RPE1 cells transiently transfected with indicated siRNA were cultured in serum-starved medium for 48 hrs. Cells were immunostained as described in Figure 4C. (A) DNA was stained with Hoechst (blue). Scale bar, 5 μm. (B) The percentage of cells with SMO-negative centrosome (cilia - & SMO -), SMO-positive centrosome (cilia - & SMO +), or SMO-positive cilia (cilia + & SMO +) was determined. Average of three independent experiments is shown. >100 cells were scored for each experiment. **(C, D)** Indicated cells were cultured in serum-starved medium for 3 (T3) or 6 (T6) hrs and immunostained with anti-glutamylated tubulin (green) and anti-Myo-Va (red) antibodies. (C) Representative Myo-Va-positive or -negative centrosome in RPE1 cells. DNA was stained with Hoechst (blue). Scale bar, 2.5 μm. (D) The percentage of cells with Myo-Va-dot at non-ciliated centrosome was determined. Average of three independent experiments is shown. >100 cells were scored for each experiment. **(E)** RPE1 cells transiently transfected with indicated siRNA were cultured and immunostained as described in Figure 5C. The percentage of cells with Myo-Va-dot at non-ciliated centrosome was determined. Average of three independent experiments is shown. >100 cells were scored for each experiment. **(F)** RPE1 cells transfected with plasmids expressing EGFP and indicated proteins were cultured and immunostained as described in Figure 5C. The percentage of cells with Myo-Va-dot at non-ciliated centrosome was determined. Average of three to four independent experiments is shown. >100 cells were scored for each experiment. **(G)** RPE1 or Rabl3-1 cells transfected with plasmids expressing EGFP and RAB11A/ Q70L were cultured and immunostained as described in Figure 5C. The percentage of cells with Myo-Va-dot at non-ciliated centrosome was determined. Average of four independent experiments is shown. >100 cells were scored for each experiment. **(B-G)** Error bars represent SD. *: *p* < 0.05, **: *p* < 0.01 compared with RPE1 (D), siLuc (E), Mock (F), or RPE1 + RAB11A/QL (G). #: *p* < 0.05, ##: *p* < 0.01 compared with siRAB8A + B (B).

We further examined CV formation by using an anti-Myo-Va antibody, which is known to mark both initial CV without GFP-SMO (Myo-Va^+^/GFP-SMO^-^ CV) and subsequent CV (Myo-Va^+^/GFP-SMO^+^ CV) (Wu et al., 2018). As expected, single Myo-Va-positive dots were detected adjacent to the centriole, and some of them grossly overlapped with GFP-SMO-dots when RPE1 cells were induced to quiescence for 6 h (Figure 5C, S3). The number of Myo-Va^+^ CV was significantly lower in RABL3-mutated cells than that in WT cells at 6 h of starvation, whereas it was equivalently observed in both cells after 3 h of starvation (Figure 5D). These results suggest that RABL3 is dispensable for initial Myo-Va^+^/GFP-SMO^-^ CV formation but is required for later Myo-Va^+^/GFP-SMO^+^ CV formation (Figure S4). A similar phenotype was also induced by the silencing of RAB11 (Figure 5E). Conversely, ectopic expression of WT or active mutants, but not negative mutants of RABL3 or RAB11A, enhanced Myo-Va^+^ CV formation (Figure 5F). Moreover, increased Myo-Va^+^ CV by overexpression of RAB11A/QL was partly but significantly inhibited in RABL3-mutated cells (Figure 5G). Collectively, these results suggest that the RAB11-RABL3 axis positively controls CV formation during ciliogenesis in human cells.

## Discussion

Of six RABL proteins, five (2A, 2B, 3, 4, and 5) are similar to typical small GTPases in size (185–266 aa in humans), in contrast RABL6 (730 aa in humans) is a relatively large protein. Based on our findings, it is possible to suggest that all small GTPase-type RABL proteins play cilia-related functions in human cells. While other small RABL proteins localize to the basal body and/or the ciliary axoneme, RABL3 shows a wider distribution throughout cells. Since RABL3 has been shown to pertain to multiple cellular events, including KRAS signaling, lymphopoiesis, and cancer cells proliferation and migration (An et al., 2017; Ge et al., 2019; Li et al., 2010; Ma et al., 2021; Nissim et al., 2019; Pan et al., 2017; Usman et al., 2020; Xu et al., 2021; Zhang et al., 2016; Zhong et al., 2020), RABL3 may be located broadly to play distinct roles depending on target molecules, interacting proteins, and/or cell types.

The RABL3-RAB11 interaction that we have demonstrated in this study is analogous to other RABL-small GTPase associations in that RABL4/IFT27 and RABL5/IFT22 bind to the small GTPase ARL6/BBS3, and these interactions occur during ciliary events. GTP-bound RABL4 associates with nucleotide-free or GDP-bound ARL6 in human cells (Liew et al., 2014). Both RABL4 and RABL5 stabilize ARL6 (Liew et al., 2014; Xue et al., 2020). Interestingly, RABL4 has been reported to be structurally akin to RAB11 (Bhogaraju et al., 2011). Therefore, our findings and previous reports indicate that similar GTP-bound small GTPases (RAB11 and RABL4) stabilize GDP-bound or nucleotide-free small GTPases (RABL3 and ARL6, respectively), both of which are related to ciliary functions in human cells. However, immunoblotting and immunostaining for RABL3 in RAB11-depleted cells did not detect a decrease in RABL3 expression (Figure 1F, data not shown). This is likely because RAB11 stabilizes RABL3 in a specific location and/or time-point during ciliogenesis. Our immunoprecipitation and subcellular localization analyses suggest that the chaperone effect of RAB11 could contribute to the recruitment of GDP-bound or nucleotide-free RABL3 to the centrosome. As RABL3/WT but not RABL3/S20N rescued de-ciliation in RABL3-mutated cells, GTP-binding is required for RABL3 function during ciliogenesis. Therefore, we hypothesize that GDP on RABL3 is probably replaced with GTP by GEF(s) around the centrosome, and then the GTP-bound RABL3 promotes CV formation through effector protein(s). Future work should identify the GEF(s) and effector(s) of RABL3 during ciliogenesis.

Presenting study may have uncovered a previously unidentified step in the CV formation pathway in human cells. RABL3 is dispensable for initial CV formation but is required for the following step, denoted “CV maturation” (Figure S4). Once the initial CV is assembled form DAVs, RABL3 is required for the subsequent “CV maturation” step in which several ciliary membrane proteins such as GFP-SMO are probably loaded to the CV. RAB11 is also involved in CV maturation through its association with RABL3. This hypothesis agrees with a previous study in which initial Myo-Va^+^ CV formation was unaffected by depletion or overexpression of RAB11 (Wu et al., 2018).

After CV maturation, RAB8 promotes the CV extension, which is also activated by RAB11 via Rabin8 (Lu et al., 2015). In the present model, RAB11 in the GTP-bound state controls continuous CM formation processes through its association with RABL3 or Rabin8-RAB8. A recent study showed that serum deprivation triggers primary cilia formation through PI3K/Akt and downstream RAB11 (Walia et al., 2019). Therefore, it is feasible that RAB11 regulates multiple steps in early ciliogenesis, including RABL3-dependent CV maturation. It will be of interest to investigate whether serum deprivation and/or the PI3K/Akt axis impact the RABL3-dependent CV maturation step. Furthermore, as RABL3-mutated zebrafish exhibited developmental phenotypes reminiscent of cilia-loss mutants (Nissim et al., 2019), future genetic studies may lead to the identification of unknown mutations in *Rabl3*, which cause cilia-related diseases.

## Materials & Methods

### Cell culture

HEK293T cells (from B. D. Dynlacht) were grown in DMEM (Nacalai tesque) supplemented with 10% Calf Serum (Thermo Fisher Scientific) and 100 units/ml penicillin and 100 μg/ml streptomycin (P/S) (Nacalai tesque). hTert-RPE1 (RPE1) (from B. D. Dynlacht), GFP-SMO-RPE1 (generated in this study), and GFP-Rabin8-RPE1 (from P. K. Jackson) were grown in DMEM supplemented with 10% Fetal Bovine Serum (FBS) (Biosera) and P/S.

### Antibodies

Antibodies used in this study include rabbit anti-RABL3 (1:500 (IF), 1:1000 (WB), Abcam, ab196024), mouse anti-glutamylated tubulin (GT335) (1:1000 (IF), Adipogen, AG-20B-0020), rabbit anti-ARL13B (1:1000 (IF), Proteintech, 17711-1-AP), rabbit anti-GM130 (1:1000 (IF), BD biosciences, #610822), mouse anti-RAB11 (1:100 (IF), 1:1000 (WB), Millipore, 05-853), rabbit anti-RAB8A (1:1000 (WB), Proteintech, 55296-1-AP), mouse anti-Centrin (1:1000 (IF), Millipore, 04-1624), rabbit anti-CP110 (1:1000 (IF), from B. D. Dynlacht) (Chen et al., 2002b), goat anti-CEP164 (1:200 (IF), Santa Cruz, sc-240226), rabbit anti-Myosin-Va (1:500 (IF), Novus, NBP1-92156), mouse anti-β-Actin (1:1000 (WB), Santa Cruz, sc-47778), rabbit anti-FLAG (1:1000 (WB), Sigma Aldrich, F7425), rabbit anti-Myc (1:1000 (WB), MBL, #562), rabbit anti-GST (1:1000 (WB), MBL, PM013), and rabbit anti-GFP (1:1000 (WB), Santa Cruz, sc-9996).

### Plasmids

To generate Flag-RABL3 or Myc-RABL3, human RABL3 fragment encoding residue 1-236 was amplified by PCR using forward primer 5’-AAGAATTCATGGCGTCCCTGGATCGGGT-3’ and reverse primer 5’-AAGTCGACTCAGTCATAATGAAGGCTCTT-3’, and sub-cloned into pCMV5-Flag or pCMV5-Myc (from K. Kontani and T. Katada). RABL3/S20N plasmid was made by PCR-based mutagenesis using forward primer 5’-AATTCGTTAGTCCATCTCCT-3’ and reverse primer 5’-TTTCCCAACACCTGAGTCTC-3’. pEGFP-C2-hRAB11A was obtained from K. Kontani and T. Katada. RAB11A/S25N and Q70L constructs were made by PCR-based mutagenesis using forward primer 5’-AATAATCTCCTGTCTCGATTTAC-3’ and reverse primer 5’-CTTTCCAACACCAGAATCTC-3’, and forward primer 5’-CTAGAGCGATATCGAGCTAT-3’ and reverse primer 5’-CCCTGCTGTGTCCCATATCT-3’, respectively. To generate Myc-RAB11A, RAB11A/S25N, or RAB11A/Q70L, hRAB11A fragments were sub-cloned into pCMV5-Myc. To generate recombinant GST-RABL3 and GST-RAB11A, hRABL3 and hRAB11A fragments were sub-cloned into pGEX6P1. To make PX459-hRABL3, annealed oligo (5’-CACCGATGACCAACGACGCAAGTTT-3’ and 5’-AAACAAACTTGCGTCGTTGGTCATC-3’) was inserted into PX459 (pSpCas9(BB)-2A-PuroV2.0) (Addgene) (Ran et al., 2013). The plasmid expressing Flag-RABL2B was previously described (Dateyama et al., 2019). pGEX6P1-GFP nanobody was obtained from Y. Katoh and K. Nakayama (Katoh et al., 2015). pEGFP-mSmo was obtained from Addgene (Chen et al., 2002a).

Plasmid transfection into HEK293T cells was performed using PEI Max (Polysciences) according to the manufacturer’s instruction. Plasmid transfection into RPE1 cells was performed using ViaFect (Promega) according to the manufacturer’s instruction.

### RNAi

siRNA oligos used in this study were siRABL3#1 (Ambion, s49874), siRABL3#2 (Ambion, s57630), siRAB8A (5’-GACAAGUUUCCAAGGAACGtt-3’, Sigma Aldrich), siRAB8B (5’-GACAAGUGUCAAAAGAAAGtt-3’, Sigma Aldrich), siRAB11A (5’-UGUCAGACAGACGCGAAAAtt-3’, Sigma Aldrich), siRAB11B (5’-GCACCUGACCUAUGAGAACtt-3’, Sigma Aldrich). The siRNA for luciferase (siLuc) was described previously (Kobayashi et al., 2017). For RNAi, 2 × 10^4^ RPE1 cells were seeded in 24-well plate and cultured for 24 hrs. After transfection of 20 pmol siRNA using Lipofectamine RNAiMAX (Invitrogen), cells were cultured in normal medium for 24 hrs and subsequently incubated in serum starved medium.

### Immunoprecipitation and Western blotting

Cells were lysed with lysis buffer (50 mM Hepes-NaOH pH 7.5, 150 mM NaCl, 5 mM MgCl_2_, 0.5% NP-40, 1 mM DTT, 0.5 mM PMSF, 2 μg/ml leupeptin, and 10% Glycerol) at 4 °C for 30 minutes. For immunoprecipitation, 1 mg of the resulting supernatant after centrifugation was incubated with anti-Flag agarose beads (Sigma Aldrich) or GFP-nanobody at 4 °C for 2 hrs. The resin was washed with lysis buffer, and the bound polypeptides were analyzed by SDS-PAGE and immunoblotting. 10 μg of lysate was loaded in the input (IN) lane.

### Immunofluorescence microscopy

Immunofluorescence microscopy was performed as described previously (Kobayashi et al., 2017). Quantification analysis of fluorescence was performed using ImageJ (Kobayashi et al., 2020).

### Generation of RABL3-mutated RPE1 cells

PX459-hRABL3 plasmid was transfected into RPE1 cells using Lipofectamine 2000 (Invitrogen). After 24 hrs, cells were cultured in medium with 10 μg/ml puromycin (Nacalai tesque) for 3 days and singly plated into 96-well plates. Genome DNA was extracted from survival cells using QuickExtract DNA Solution 1.0 (epicentre), and mutations were determined by sequencing against amplified PCR products including guide RNA target sequence. Primers for PCR amplification and sequencing are listed in Table S1.

### Generation of GFP-SMO-RPE1 cells

To generate GFP-SMO-expressed cells, RPE1 cells were transfected with pEGFP-mSMO plasmid and subsequently cultured in medium with 1 mg/ml G418 (Nacalai tesque) for 12 days. Isolated colonies were examined by immunofluorescence.

### Recombinant proteins

*E. coli* BL21(DE3) pLysS cells (TOYOBO) harboring pGEX6P1, pGEX6P1-RABL3 or pGEX6P1-RAB11A were cultured in LB/2% Glucose medium at 37 °C until OD_600_ = 0.3. After 0.2 mM isopropyl β-D-1-thiogalactopyranoside (IPTG, Nacalai tesque) treatment, cells were subsequently incubated at 37 °C for 2 hrs. Collected cells by centrifugation were suspended with extraction buffer (50 mM Tris-HCl (pH 8.0), 100 mM NaCl, 5 mM MgCl_2_, 1 mM DTT, 10% Glycerol, 0.5 mM PMSF, 2 μg/ml leupeptin), and solubilized with Triton-X 100 with final concentration of 0.5% after sonication. The supernatant after centrifugation at 17,000 g at 4 °C for 1 hr was incubated with Glutathione Sepharose 4B (GE healthcare) at 4 °C for 3 hrs. After washing with Wash Buffer (50 mM Tris-HCl (pH 8.0), 100 mM NaCl, 5 mM MgCl_2_, 1 mM DTT, 10% Glycerol, 0.5% Triton-X100), the resin was incubated with Wash Buffer containing 15 mM Glutathione (Nacalai tesque) at 4 °C for 20 min to obtain GST or GST-RABL3.

The resin after washing was incubated with Wash Buffer containing PreScission Protease (GE healthcare) to elute RAB11A by cleavage at 4 °C for 16 hrs. The eluate containing recombinant proteins was passed through a PD-10 column (GE healthcare) to exchange buffer with Store Buffer (50 mM Tris-HCl (pH 7.5), 100 mM NaCl, 1 mM MgCl_2_, 1 mM DTT, 10% Glycerol, 0.1% Triton-X100). For Turbidity assay, proteins were solubilized in Store Buffer-2 (50 mM Tris-HCl (pH 7.5), 100 mM NaCl, 1 mM MgCl_2_, 1 mM DTT).

### GFP-nanobody

*E. coli* BL21(DE3) pLysS cells harboring pGEX6P1-GFP nanobody were cultured in LB/2% Glucose medium at 37 °C until OD_600_ = 0.5. After 0.1 mM IPTG treatment, cells were subsequently incubated at 20 °C for 20 hrs. Collected cells by centrifugation were lysed and purified using Glutathione-Sepharose 4B as described previously (Katoh et al., 2015).

### *In vitro* binding assay

0.5 μM RABL3 and 0.05 μM GST or GST-RAB11 were mixed with Glutathione-Sepharose 4B in the presence of 10 μM GTPγS (Sigma Aldrich) or 10 μM GDP (Sigma Aldrich) in Store Buffer. The mixture was incubated at 25 °C for 1 hr, and then rotated at 4 °C for 2 hrs. The resin was washed with Store Buffer, and the bound polypeptides were analyzed by SDS-PAGE and immunoblotting.

### Turbidity assay

20 μM RABL3 and 20 μM GST or GST-RAB11 were mixed on ice in Store Buffer-2. The mixture was incubated at 37 °C, and optical density (OD) was measured at 350 nm using Nanodrop 2000c (Thermo Scientific) every 3 mins.

### Identification of RABL3-interacting proteins

The procedures were basically performed as described previously (Kobayashi et al., 2011). Briefly, Flag-RABL3 or Flag-RABL2B was expressed in HEK293T cells and immunoprecipitated with anti-Flag agarose beads at 4 °C for 3 hrs. Bound proteins were eluted with Flag peptide (Sigma Aldrich) for 30 min, and the resultant eluates were TCA precipitated and analyzed by SDS-PAGE. After Coomassie staining, the gel was subjected to mass spectrometric analysis performed at NAIST mass spectrometric laboratory.

### Statistical Analysis

The statistical significance of the difference was determined using two-tailed Student’s *t*-test (except for Fig. 5B) or *x*-squared test (Fig. 5B). Differences were considered significant when *p* < 0.05. **, *p* < 0.01; *, *p* < 0.05, ##, *p* < 0.01; #, *p* < 0.05.

## Acknowledgements

We thank B. D. Dynlacht (New York University) for an anti-CP110 antibody, hTert RPE1, and HEK293T cells; P. K. Jackson (Stanford University) for GFP-Rabin8-RPE1 cells; Y. Katoh and K. Nakayama (Kyoto University) for pGEX6P1-GFP nanobody; T. Katada and K. Kontani (University of Tokyo) for pCMV5-Flag, pCMV5-Myc, and pEGFP-C2-hRAB11A. We thank Y. Fukao and R. Kurata (NAIST) for mass spectrometric analysis.

T. K. was supported by grants from JSPS KAKENHI (15H01215, 15K07931, 18K06627, 21K06528), The Kurata Memorial Hitachi Science and Technology Foundation, Takeda Science Foundation, Daiichi Sankyo Foundation of Life Science, Sagawa Foundation for Promotion of Cancer Research, Mochida Memorial Foundation for Medical and Pharmaceutical Research and Foundation for Nara Institute of Science and Technology.

