## Supplemental Figures for "Atypical small GTPase RABL3 interacts with RAB11 to regulate early ciliogenesis in human cells"

A)

**Rabl3-1 (exon 4)**

```
WT       TCCTCCCAAAACTTGCGTCGTTGGTCATTGGAA
Allele_A TCCTCCC----CTTGCGTCGTTGGTCATTGGAA
Allele_B TCCTCCCA---CTTGCGTCGTTGGTCATTGGAA
```

**Rabl3-2 (exon 4)**

```
WT       TCCTCCCAAAA CTTGCGTCGTTGGTCATTGGAA
Allele_A TCCTCCCAAAA --TGCGTCGTTGGTCATTGGAA
Allele_B TCCTCCCAAAAACTTGCGTCGTTGGTCATTGGAA
```

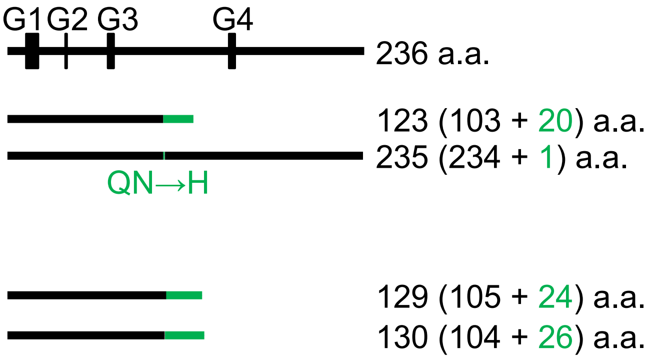


B)

```
Homo Sapiens              91 IIFVHDLTNKKSSQNLRRWS 110
Pan Troglodytes           91 IIFVHDLTNKKSSQNLRRWS 110
Canis lupus familiaris    91 IILVHDLTNKKSSQNLYRWS 110
Bos Taurus                91 IILVHDLTNKKSSQNLYRWS 110
Mus musculus              91 IILVHDLTNKKSSQNLYRWS 110
Rattus norvegicus         91 IILVHDLTNKKSSQNLYRWS 110
Gallus gallus             91 IILVHDLTNKKSSQNLYRWS 110
Danio rerio               91 IILVHDLTNKKSSQNLYRWS 110
```

**Figure S1**

**Mutations in RABL3-mutated RPE1 cells**

**(A)** (left) Mutations of the RABL3 gene in Rabl3-1 or Rabl3-2 cells. Red sequences show the target of guide RNAs with underlined protospacer adjacent motif (PAM). (right) Conserved domains of GTP-binding proteins were shown from G1 to G4. Proteins translated from each clone. Green shows incorrect residues. **(B)** Alignment of RABL3 proteins in vertebrate including mutated di-peptide (green) in allele_B of Rabl3-1.

**Figure S1**

A)

| | Mock | RABL3 | RABL2B |
|---|---|---|---|
| **Total score** | 29351 | 62706 | 257997 |
| **RAB11 score** | 0 | 191 | 0 |

B)

```
hRAB11A     MGTRDDEYDYLFKVVLIGDSGVGKSNLLSRFTRNEFNLESKSTIGVEFATRSIQVDGKTI
hRAB11B     MGTRDDEYDYLFKVVLIGDSGVGKSNLLSRFTRNEFNLESKSTIGVEFATRSIQVDGKTI
            ************************************************************

hRAB11A     KAQIWDTAGQERYRAITSAYYRGAVGALLVYDIAKHLTYENVERWLKELRDHADSNIVIM
hRAB11B     KAQIWDTAGQERYRAITSAYYRGAVGALLVYDIAKHLTYENVERWLKELRDHADSNIVIM
            ************************************************************

hRAB11A     LVGNKSDLRHLRAVPTDEARAFAEKNGLSFIETSALDSTNVEAAFQTILTEIYRIVSQKQ
hRAB11B     LVGNKSDLRHLRAVPTDEARAFAEKNNLSFIETSALDSTNVEEAFKNILTEIYRIVSQKQ
            **************************.*************** **:.*************

hRAB11A     MSDRRENDMSPSNNVVPIHVPPTTENKP--KVQCCQNI
hRAB11B     IADRAAHDESPGNNVVDISVPPTTDGQKPNKLQCCQNL
            ::**  :* **.**** * *****:.:   *:*****:
```

**Figure S2**

**RAB11 is a putative binding partner of RABL3**

(**A**) Mascot scores of Mass spectrometry analysis. **(B)** Alignment of human RAB11A and RAB11B protein sequences. Red sequences indicate identified peptides in Mass spectrometry analysis.

**Figure S2**

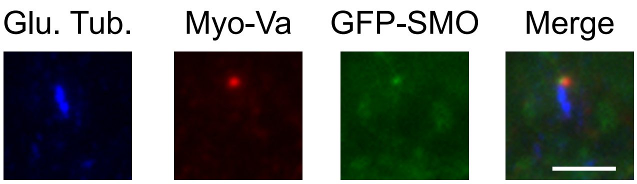


**Figure S3**

**Myo-Va dot overlaps with GFP-SMO dot in RPE1 cells**

GFP-SMO-RPE1 cells were cultured in serum-starved medium for 6 hrs. Cells were immunostained with anti-glutamylated tubulin (blue) and anti-Myo-Va (red) antibodies. Scale bar, 2.5 $\mu$m.

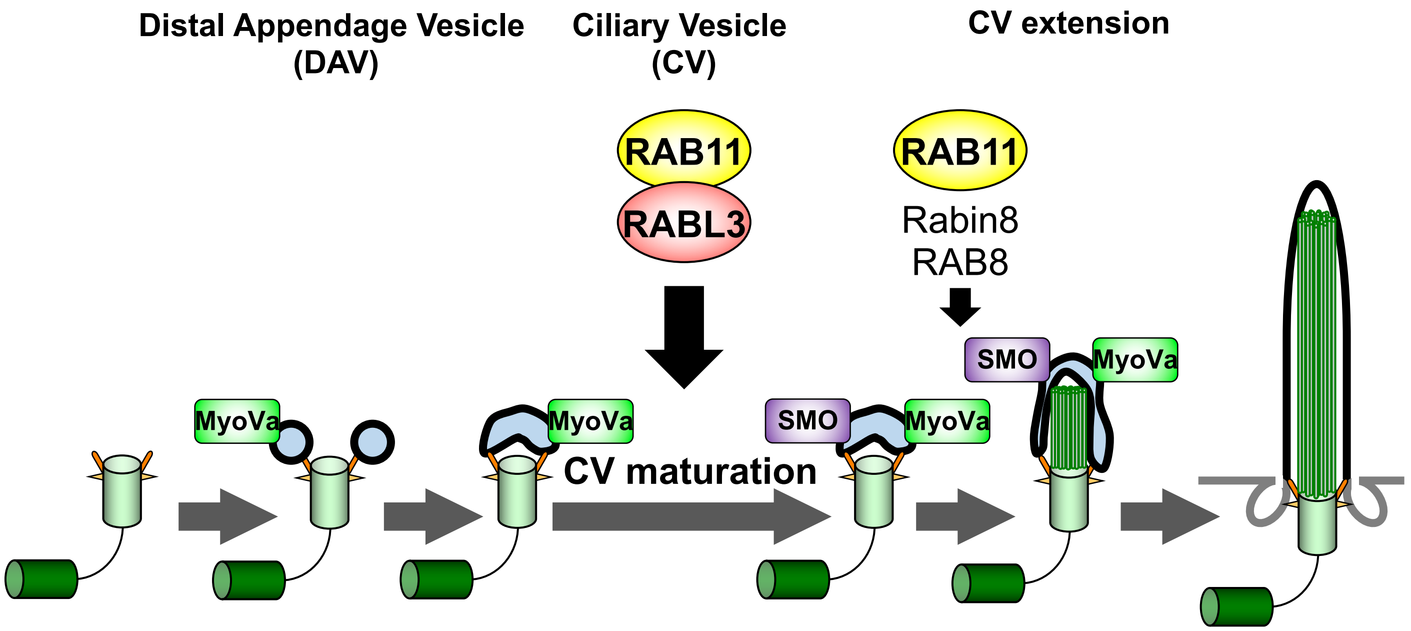


**Figure S4**

**Hypothetic model of CV formation in human cells**

RAB11-RABL3 complex mediates a CV formation step named “CV maturation” during ciliogenesis in human cells.

| Name | Sequence (5' to 3') | Use |
|---|---|---|
| hRabl3_36255_F | GTCTCTGAGTTGTTAGTCATACT | PCR |
| hRabl3_36558_R | CAAAGTGTTATTCAGGGAAGTAC | PCR |
| hRabl3_36322_F | GCTTCATTTGGCCAATTGTG | Sequencing |

**Table S1**
